# Calcium and Integrin-binding protein 2 (CIB2) controls force sensitivity of the mechanotransducer channels in cochlear outer hair cells

**DOI:** 10.1101/2023.07.09.545606

**Authors:** Isabel Aristizábal-Ramírez, Abigail K. Dragich, Arnaud P.J. Giese, K. Sofia Zuluaga-Osorio, Julie Watkins, Garett K. Davies, Shadan E. Hadi, Saima Riazuddin, Craig W. Vander Kooi, Zubair M. Ahmed, Gregory I. Frolenkov

## Abstract

Calcium and Integrin-Binding Protein 2 (CIB2) is an essential subunit of the mechano-electrical transduction (MET) complex in mammalian auditory hair cells. CIB2 binds to pore-forming subunits of the MET channel, TMC1/2 and is required for their transport and/or retention at the tips of mechanosensory stereocilia. Since genetic ablation of CIB2 results in complete loss of MET currents, the exact role of CIB2 in the MET complex remains elusive. Here, we generated a new mouse strain with deafness-causing p.R186W mutation in *Cib2* and recorded small but still measurable MET currents in the cochlear outer hair cells. We found that R186W variant causes increase of the resting open probability of MET channels, steeper MET current dependence on hair bundle deflection (I-X curve), loss of fast adaptation, and increased leftward shifts of I-X curves upon hair cell depolarization. Combined with AlphaFold2 prediction that R186W disrupts one of the multiple interacting sites between CIB2 and TMC1/2, our data suggest that CIB2 mechanically constraints TMC1/2 conformations to ensure proper force sensitivity and dynamic range of the MET channels. Using a custom piezo-driven stiff probe deflecting the hair bundles in less than 10 µs, we also found that R186W variant slows down the activation of MET channels. This phenomenon, however, is unlikely to be due to direct effect on MET channels, since we also observed R186W-evoked disruption of the electron-dense material at the tips of mechanotransducing stereocilia and the loss of membrane-shaping BAIAP2L2 protein from the same location. We concluded that R186W variant of CIB2 disrupts force sensitivity of the MET channels and force transmission to these channels.

## INTRODUCTION

Inner ear hair cells detect sound through deflection of specialized mechanosensory projections, known as stereocilia (reviewed in ^1–3^). Deflections of stereocilia result in tensioning intracellular tip links that convey mechanical force to the mechano-electrical transducer (MET) channels ^4, 5^. A mature tip link consists of a hetero-tetramer of cadherin 23 (CDH23) at the upper end and at least one of protocadherin 15 (PCDH15) isoforms at the lower end ^6–9^. In mammalian auditory hair cells, MET channels are localized at the lower end of the tip links ^10^. Therefore, the force from PCDH15 should be transmitted to the MET channels either through the plasma membrane and/or molecular partners of PCDH15. Since mammals hear at tens of kilohertz frequencies, this force transmission must be extremely fast and ensure activation of MET channels within a few microseconds. This speed is yet unachievable by piezo-driven probes commonly used in MET current recordings ^11^.

The identity of molecules forming MET machinery in mammalian hair cells is one of the hottest subjects in auditory neuroscience. Recent consensus emerges that TMC1 and TMC2 proteins form the pore of the MET channel ^12–14^. However, several potential accessory subunits of this channel have also been identified, including TMHS/LHFPL5 ^15^, TMIE ^16^, and, as we showed previously, CIB2 ^17, 18^. All of them are essential for proper assembly of the MET complex. Corresponding knockout mouse models either lack MET currents in the auditory hair cells completely (TMIE and CIB2) ^16, 18–20^, or exhibit significantly reduced MET currents (LHFPL5) ^15, 21^. *In vitro*, LHFPL5 and TMIE interact with PCDH15 ^15, 16, 22^, while TMIE and CIB2 can interact directly with TMC1 and TMC2 ^18–20^. In addition, TMC proteins can interact with PCDH15 ^23^, while TMIE could also bind to LHFPL5 ^16^. It is yet unknown how exactly all these potential interactions assemble MET machinery and which of them are present in the mature MET complex. The exact mechanism of MET channel activation is also uncertain. On one hand, bird and reptile orthologs of TMC1 truncated at C- and N-termini can form functional mechanosensory ion channels when reconstituted in artificial liposomes ^24^, suggesting that TMC-based MET channels may be activated by membrane stretch like many other mechanosensory channels. This mechanism of gating by membrane tension is also consistent with well-established lipid effects on the MET currents ^25, 26^. On the other hand, there are also strong arguments for the force transmission from the tip link to the MET channel via accessory protein LHFPL5 ^27^.

As an auxiliary subunit of the MET complex, CIB2 is particularly interesting because it may be responsible for at least some of well-known effects of Ca^2+^ on hair cell transduction. We and others previously identified multiple variants in *CIB2* gene responsible for hearing loss in diverse populations ^17, 28–31^. Some of these deafness-causing variants disrupt CIB2 interactions with TMC1/2 *in vitro*, suggesting that CIB2 is a part of MET complex ^18^. A recent structural study ^19^ has determined how CIB proteins interact with TMCs but the exact function of CIB2 in MET complex remains elusive.

To explore the role of CIB2 in MET function, we have generated a knock-in mouse strain (*Cib2^R186W^*), which recapitulates one of the variants of human deafness that is less deleterious for interaction of CIB2 with TMC1/2 ^18^. Although these mice are profoundly deaf, their outer hair cells (OHCs) exhibit decreased but still detectable MET currents. The most prominent effects of the R186W variant on these currents were increased sensitivity of the MET channels to the hair bundle deflection (steeper I-X curves), increased resting open channel probability (P_open_), and exaggerated leftward shifts of the I-X curves upon cell depolarization. All these effects indicate a larger sensitivity of *Cib2^R186W/R186W^* MET channels to the gating force. AlphaFold2 modeling confirmed that, indeed, p.R186W mutation may disrupt one of the multiple interactions between CIB2 and TMC1/2, thereby releasing mechanical constraints to the MET channel. In addition, using a very fast microfabricated piezo probe, we were able to deflect OHC bundles fast enough to resolve the time constant of MET channel activation. We found that R186W variant significantly slows down MET channel activation in OHCs. However, the latter effect may be due to delayed force transmission to the MET channels - we found profound ultrastructural changes at the tips of transducing stereocilia in *Cib2^R186W/R186W^* hair cells as well as the loss of BAR-domain protein BAIAP2L2 that may affect the mechanical properties of the plasma membrane in the vicinity of the MET channels.

## RESULTS

### *CIB2^R186W^*: a mouse model to decipher the role of CIB2 in the MET complex

Profound deafness caused by mutations in *CIB2* in humans and by genetic deletion of *Cib2* in mice, complete loss of the MET currents in CIB2-deficient auditory hair cells, and strong binding of CIB2 to TMC1 and TMC2 *in vitro* have established CIB2 as an integral accessory subunit of the MET complex ^17–19^. However, since both genetic deletion of *Cib2* and point mutations disrupting its interaction with TMC1/2 result in complete loss of the MET currents in the auditory hair cells ^18, 19^, the exact role of this protein within the MET complex remains elusive. Out of six deafness-causing human variants of *CIB2*, three (p.E64D, p.F91S, and p.C99W) disrupt the interactions of CIB2 with TMC1 *in vitro* (**Figure 1A, red**), while the other three (p.R66W, p.I123T, and p.R186W) retain at least some level of these interactions (**Figure 1A, green**) ^18^. Since the p.R186W variant does not affect CIB2 trafficking to the hair cell stereocilia ^28^, we reasoned that a detail examination of the MET currents in the auditory hair cells of mice carrying R186W variant could help in deciphering the role of CIB2 in the MET-complex. Therefore, we developed a *Cib2* knock-in mouse strain (*Cib2^R186W^*) harboring p.R186W variant (**Figure 1B**).

**Figure 1.**
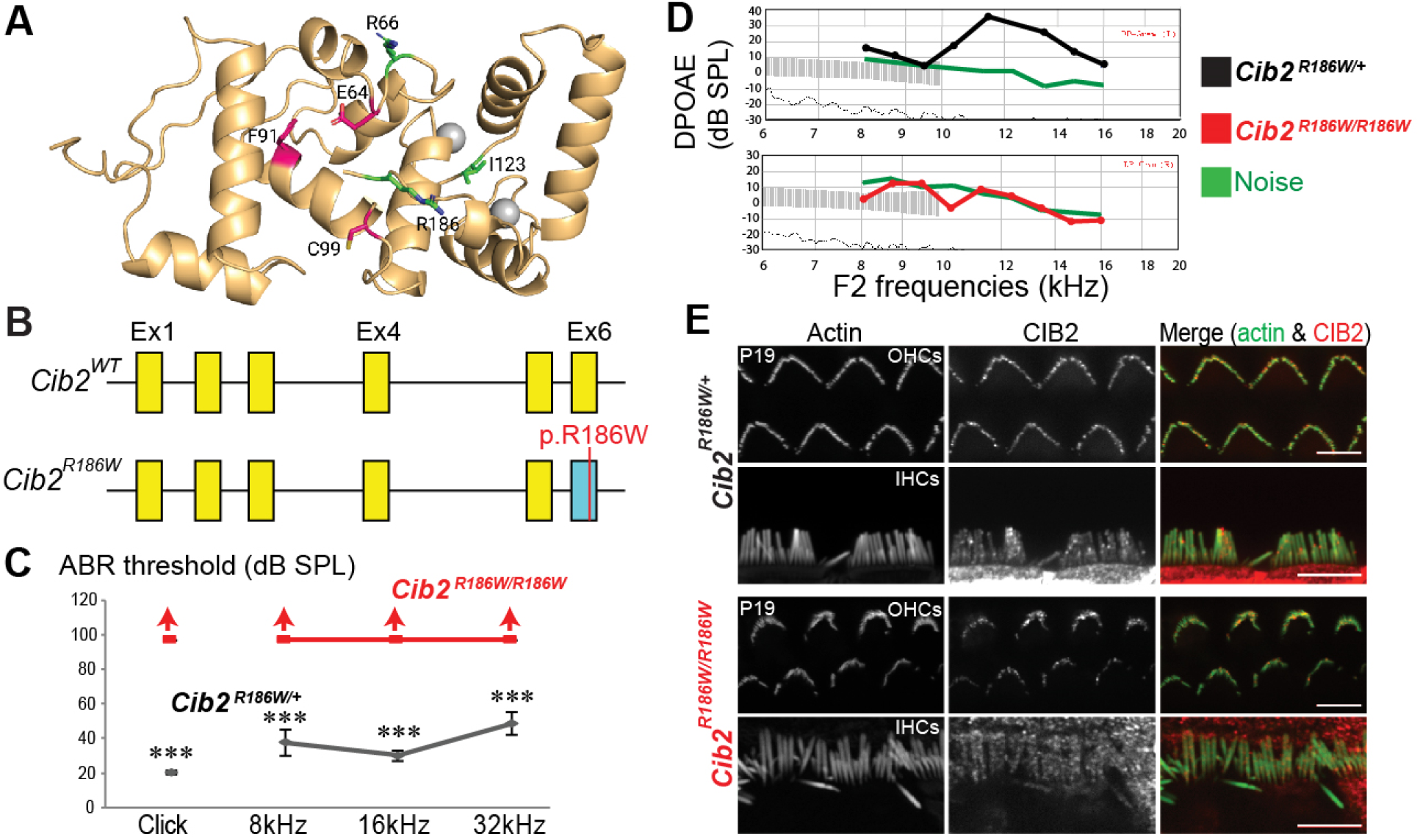
*Cib2^R186W^* mutant mice are profoundly deaf. (A) An Alphafold ^63^ model of wild type CIB2. The two conserved calcium binding sites observed in CIB1 (PDB 1XO5) and CIB3 (PDB 6WU7) are shown as grey spheres. Although in both CIB1 and CIB3, other calcium binding cites were reported but these are thought to be due to the very high calcium concentrations used for crystallization ^19^. The experimental crystal structure for CIB2 is not yet available. CIB2 residues mutated in six missense pathogenic variants associated with hearing loss are shown. Variants in red completely abolish the CIB2-TMC1 interaction in pull-down assays while variants in green retain some CIB2-TMC1 interaction ^18, 19^. (B) Structure of *Cib2^R186W^* alleles generated by CRISPR/Cas9 technology. CIB2^R186W^ protein structure is shown. (C) ABR thresholds (dB SPL) to broadband clicks and tone-pips with frequencies of 8 kHz, 16 kHz, and 32 kHz in *Cib2^R186W/+^* (black) and *Cib2^R186W^*^/*R186W*^ (red) mice at P21. (D) DPOAEs of control (black) and *Cib2^R186W^*^/*R186W*^ (red) mice at P21. Noise floor is shown in Green. All data are shown as Mean ± SEM (\*\*\**p* < 0.001). (E) Maximum intensity projections of confocal Z-stacks of the medial turns of *Cib2^+/+^* and *Cib2^R186W/R186W^*organs of Corti immunostained with CIB2 antibody (red) at P19 and counterstained with phalloidin (green). Scale bars: 15 μm.

First, we characterized the hearing function in *Cib2^R186W^*mutant mice. Auditory brainstem response (ABR) evaluation revealed normal waveforms and thresholds in mice heterozygous of R186W variant (**Figure 1C**). In contrast, *Cib2^R186W^* homozygous mutant mice had no response to either click or tone burst stimuli of 100 dB sound pressure level (SPL) indicating profound deafness (**Figure 1C**). Similarly, distortion product otoacoustic emission (DPOAE) a by-product of cochlear amplification that depends on the integrity of OHCs, was indiscernible from the noise floor in the mutant mice at all tested frequencies (**Figure 1D**). However, *Cib2^R186W/R186W^* mice did not display any obvious indications of vestibular dysfunction, such as circling, hyperactivity or head bobbing, indicating predominantly auditory phenotype. Scanning Electron Microscopy (SEM) did not show profound degeneration of the auditory hair cell bundles at P17 in *Cib2^R186W/R186W^* mice. However, the OHC bundles exhibited the same characteristic abnormalities that we previously reported for another *Cib2* mutant strain (*Cib2^F91S^*) ^18^ – the “horseshow” shape of the hair bundle, retraction of many stereocilia in the 3^rd^ row, and nearly equal heights of the remaining stereocilia in all three rows of an OHC bundle (**Supplemental Fig. 1A**). We also did not observe a widespread auditory hair cells loss at P21, except the loss of OHCs at the base of the cochlea (**Supplemental Fig. 1B**).

To determine whether the auditory deficit in *Cib2^R186W/R186W^* mice is due to disruption of CIB2 function or its localization, we performed immunofluorescent labeling with CIB2 antibodies that we previously validated in *Cib2* knockout mice ^18^. Consistent with the idea on functional deficiency of the R186W variant of CIB2 within the MET complex, we observed a very similar and distinct CIB2 labelling in stereocilia of both inner hair cells (IHCs) and OHCs and in both *Cib2^R186W/+^* and *Cib2^R186W/R186W^*mice (**Figure 1E**). We hypothesized that R186W variant may affect MET currents in the auditory hair cells due to the changes in CIB2 interaction with TMC1/2 and/or other molecular partners.

### Intact tip links and overgrown transducing stereocilia in the auditory hair cells of *CIB2^R186W/R186W^* mice

The first signs of abnormalities in *Cib2^R186W/R186W^*stereocilia bundles were observed in young postnatal mice already at P6-P8 (**Figure 2**). At this age, the OHC bundles begin to acquire horseshoe shapes and had already somewhat disrupted staircase arrangement of stereocilia rows, while the overall structure of IHC bundles appears relatively normal **(Figure 2A-C)**. However, the 2^nd^ row stereocilia in *Cib2^R186W/R186W^* IHCs already exhibit abnormally elongated tips (**Figure 2B,D**) and the overall length of transducing stereocilia in both 2^nd^ and 3^rd^ rows were increased (**Figure 2E, left**). In the *Cib2^R186W/R186W^* OHCs, the remaining (not retracted) transducing stereocilia are also overgrown (**Figure 2E, right**).

**Figure 2.**
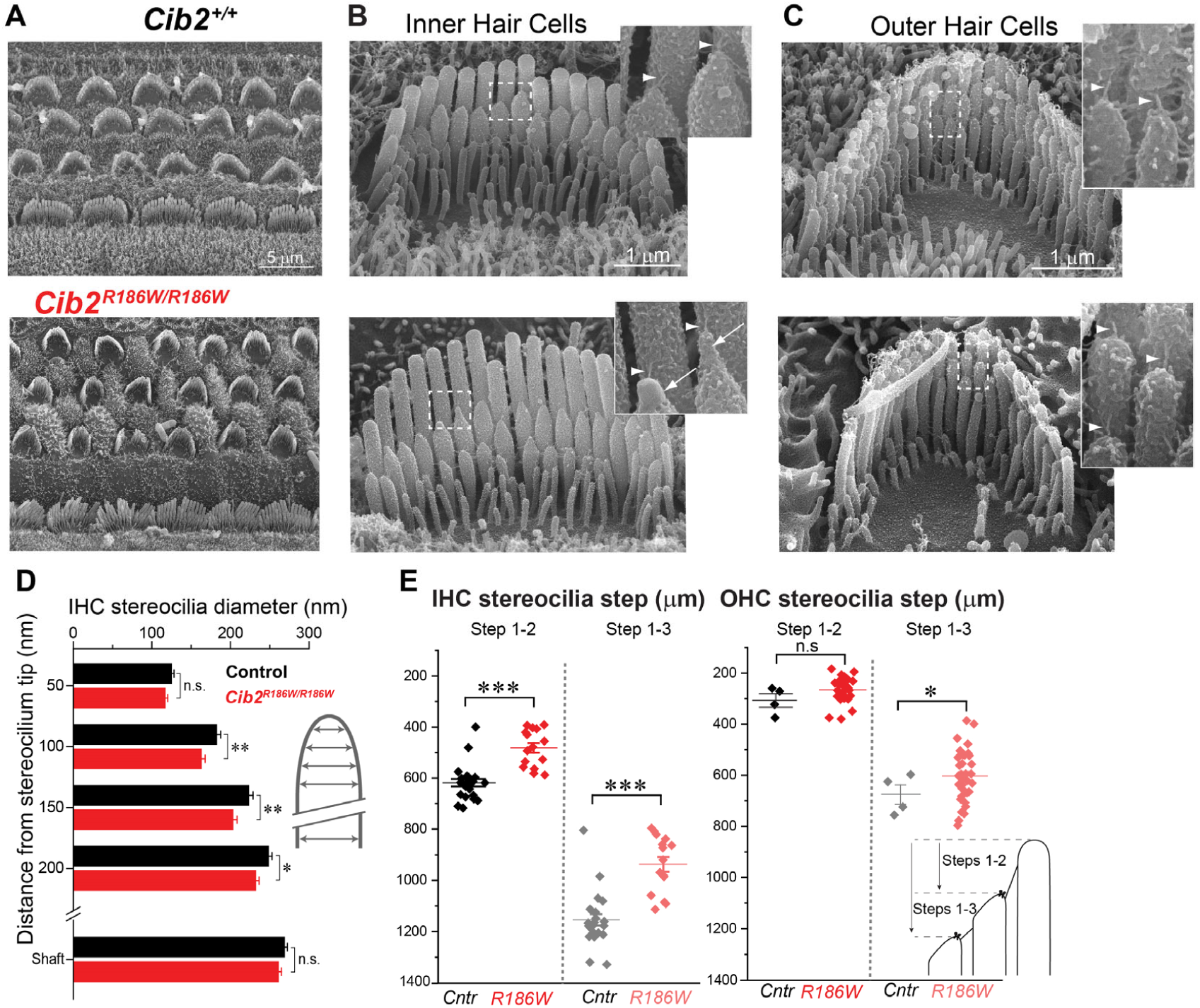
R1186W variant causes characteristic changes of the staircase organization of stereocilia in the auditory hair cells without affecting tip links. (A) Scanning electron microscopy (SEM) images of the organs of Corti explants from young (P6-P8) postnatal wild type (*Cib2^+/+^*, top) and *Cib2^R186W/R186W^* (bottom) mice in the middle of the apical turn of the cochlea. Note a more rounded “U-shape” of the OHC bundles in R186W mutants. (B-C) Representative SEM images of IHC (B) and OHC (C) bundles from the same samples. Insets show magnified views of the areas indicated by dashed rectangles. Identical magnification is used for images and insets in the left and right panels. Arrowheads indicate intact tip links in IHC and OHC. Arrows point to the abnormalities of the shape of stereocilia tips in *Cib2^R186W/R186W^*IHCs. (D) “Overtented” tips of IHC stereocilia in *Cib2^R186W/R186W^* mice. Diameters of the second row stereocilia were measured every 50 nm down from the tip of each stereocilium. (E) “Step distances” between the tip of the first row and the tip of the second row (steps 1-2) or the tip of the third row (steps1-3) in *Cib2^R186W/R186W^* and control IHCs and OHCs (see an explanatory cartoon in an inset). Completely retracted 3^rd^ row stereocilia in OHCs were not included in the analysis to quantify over-elongation of the remaining 3^rd^ row stereocilia. Each data point represents individual stereocilium. Error bars show mean±SD. Asterisks indicate significance: *, p<0.05; **, p< 0.005; ***, p<0.0005 (Student *t*-test of independent variables). Age of the cells: P6-P8.

Despite these changes in the hair bundle morphology, both IHCs and OHCs in the young postnatal *Cib2^R186W/R186W^* mice have prominent tip links (**Figure 2B,C, arrowheads**). This was confirmed by tip link count, were there was no differences between wild type and *Cib2^R186W/R186W^* cells in the percentage of stereocilia pairs possessing tip links per hair bundle (Mean±SE): 76.8±2.7%, n=5, wild type OHCs; 73.5±1.8%, n=9, *Cib2^R186W/R186W^* OHCs; 78.4±1.2%, n=4, wild type IHCs; 74.8±1.2%, n=7, *Cib2^R186W/R186W^* IHCs (differences are not significant; Student *t*-test of independent variables).

### *Cib2^R186W/R186W^* OHCs have fewer MET channels that are, however, more sensitive to hair bundle deflections

Next, we explored whether *Cib2^R186W/R186W^* OHCs are mechanosensitive. We used a piezo-driven rigid probe to deflect stereocilia bundles and a conventional whole-cell patch clamp to record the resulting MET responses (**Figure 3A, inset**). In contrast to p.F91S mutation that results in no measurable MET currents ^18^, positive deflections of the hair bundle in *Cib2^R186W/R186W^* OHCs caused small but detectable MET currents (**Figure 3A**), with an average amplitude significantly smaller than in wild type OHCs (**Figure 3B**). Consistent with the normal hearing in the adult *Cib2^R186W/+^* mice **(Figures 1C,D)**, the amplitudes of MET currents in *Cib2^R186W/+^* OHCs were not statistically different from that in wild type OHCs (**Figure 3B**). Normalized relationships between peak MET current and hair bundle deflection (known as “I-X” curves) revealed that the MET channels in *Cib2^R186W/R186W^* OHCs are more sensitive to hair bundle deflections (have steeper I-X curve) compared to *Cib2^R186W/+^*and wild type OHCs **(Figure 3C)**. We quantified this phenomenon by calculating the “width of I-X curve” between 25% and 75% of the total MET current and found highly significant difference between wild type and *Cib2^R186W/R186W^*OHCs (**Figure 3D**).

**Figure 3:**
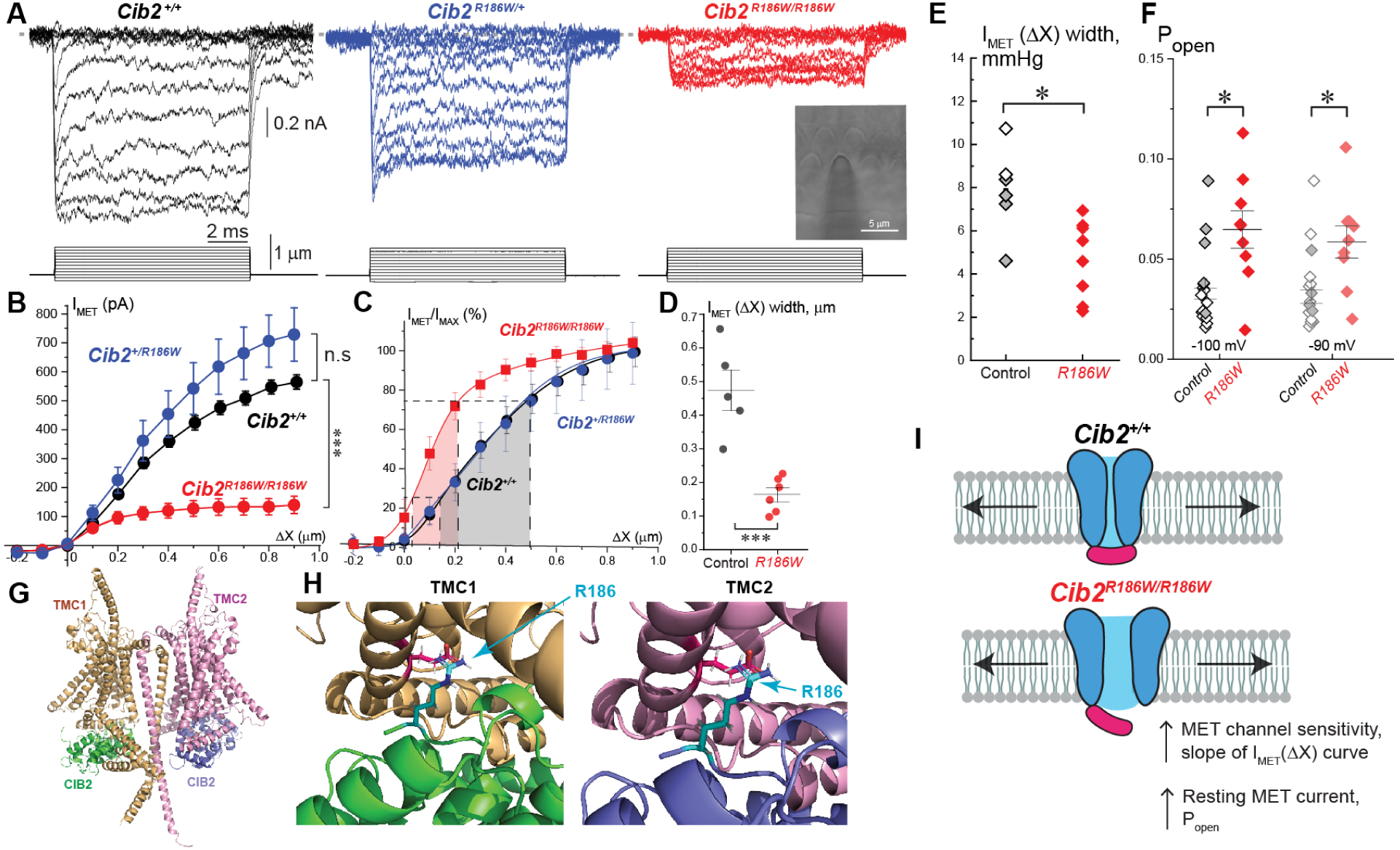
*Cib2^R186W/R186W^* OHCs have reduced MET currents but increased sensitivity of MET channels to an external force. (A) Representative traces of the MET currents (top) evoked by the graded step-like deflections (bottom) of stereocilia bundle with the stiff-probe in *Cib2^+/+^*(left), *Cib2^R186W/+^* (middle) and *Cib2^R186W/R186W^*(right) OHCs. Inset on the right panel shows bright-field image of the organ of Corti explant with the stiff probe deflecting OHC bundle and a recording patch clamp pipette underneath the focal plane (dark area on the right). (B) Relationship between the peak MET current (I_MET_) and hair bundle displacement (ΔX). Data are shown as mean±SE. Asterisks indicate statistical significance: ***, p<0.0005 (two-way ANOVA). (C) Relationships between normalized MET current (I_MET_/I_MAX_) and hair bundle displacement, further referred to as I-X curves. Curves were fitted with a double Boltzmann curve. Shaded regions under the curves (grey for control and light red for *Cib2^R186W/R186W^*) highlight the 25% to 75% increase in the MET channel opening that was used to quantify the steepness of the I-X curves shown in D. (D) Width of I-X curves (at 25-75% of MET channel activation) in *Cib2^+/+^* and *Cib2^R186W/R186W^* OHCs. Asterisks indicate significance: ***, p<0.0005 (Student *t*-test of independent variables). (E) Widths of I-X curves determined as in D but in an independent series of experiments with the fluid-jet deflection of the OHC bundles. To minimize cell-to-cell variability, only the cells deflected with the pipettes of 4.0 ± 0.1 µm in diameter were included in this quantification. Control group includes both *Cib2^+/+^* (open symbols) and *Cib2^R186W/+^* (filled symbols) OHCs. (F) Resting open probability of MET channels (P_open_) at negative holding potentials (-100 mV, -90 mV) from all experiments with fluid-jet deflections of stereocilia bundles. Individual data from each OHC as well as mean±SE are shown. Symbols filled with grey correspond to *Cib2^R186W/+^*cells. Asterisks indicate significance: *, p<0.05 (Student *t*-test of independent variables). Number of cells/mice: control, n=16/6 (*Cib2^+/+^*, n=12/4; *Cib2^R186W/+^*, n=4/2); *Cib2^R186W/R186W^*, n=9/2. (G) AlphaFold2 multimer prediction of CIB2 (green and blue) interaction with TMC1 (tan) and TMC2 (pink) within the heterocomplex. (H) In the two copies of CIB2, R186 (cyan) forms a salt bridge with either TMC1 E172 (left) or TMC2 E224 (right). (I) In a wildtype OHC (top), multiple CIB2-TMC1/2 interactions may mechanically constrain the MET channel, thereby setting its sensitivity to the gating force and its resting open probability. R186W variant may release those constraints and makes the channel more responsive to the tension within the MET complex (bottom). Age of the cells: P4-P6; all of them were located approximately in the middle of the apical turn of the cochlea.

However, the shape of I-X curve in the above experiments may be affected by the fact that the piezo-driven rigid probe is ineffective in deflecting all stereocilia within the hair bundle simultaneously ^32^. Therefore, we performed a separate series of experiments, where we recorded MET currents evoked by deflection of OHC bundles with a fluid-jet **(Figure 4A, inset)**. Consistent with the rigid probe data, we observed a substantial decrease of the amplitude of MET currents in *Cib2^R186W/R186W^* OHCs (**Supplemental Fig. 2B, left**). The decrease of the MET currents in *Cib2^R186W/R186W^* OHCs cannot be explained by a simple loss of tip links (**Figure 2C**) and, therefore, it must be a result of decreased number of MET channels and/or decreased single MET channel conductance. Therefore, we estimated the single MET channel conductance with non-stationary fluctuation analysis of the MET currents produced by ramp-like deflections of the OHC bundles with a fluid-jet (**Supplemental Fig. 2A**). This sort of analysis does underestimate single MET channel conductance but can be used for comparison of OHCs with different genotypes, at least in the apical OHCs ^33^. We found no differences in the estimated single MET channel conductance between control (*Cib2^+/+^* and *Cib2^R186W/+^*) and *Cib2^R186W/R186W^* OHCs at P4-P6 (**Supplemental Fig. 2B, right**) but an increased single channel conductance in the control OHCs at P7 (**Supplemental Fig. 3A**), perhaps, due to an increased number of TMC1-only MET channels in the control OHCs (see Discussion below). Therefore, we have limited any further analysis to the OHCs at the ages of P4-P6. At these ages, the same *Cib2^R186W/R186W^* OHCs have apparently normal single MET channel conductance but several-fold smaller MET currents (**Supplemental Fig. 2B**), clearly indicating fewer functional MET channels in *Cib2^R186W/R186W^* OHCs compared to the control.

**Figure 4:**
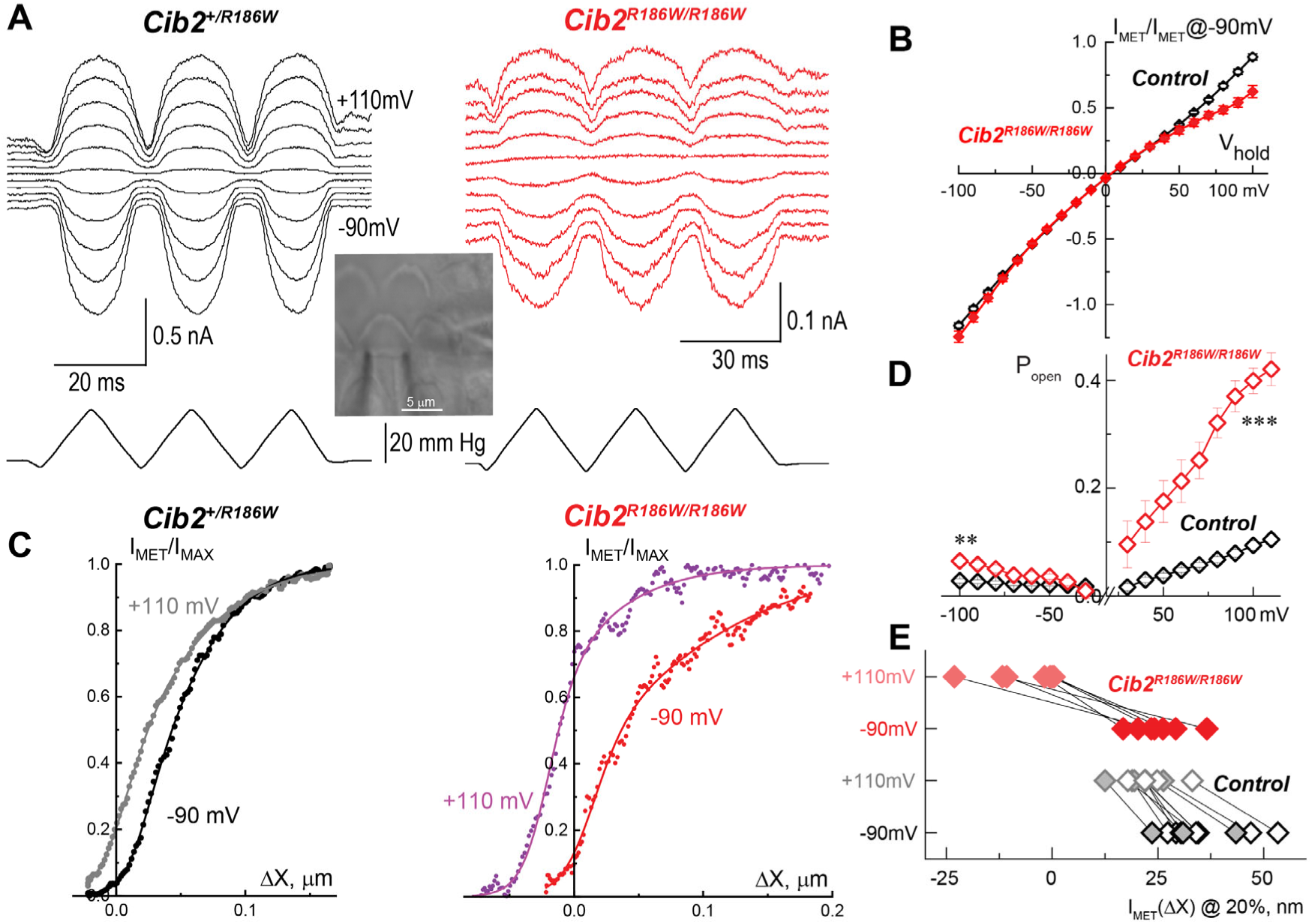
R186W variant exacerbates voltage-dependence of the MET currents. (A) Representative MET currents (top) of control (left) and *Cib2^R186W/R186W^* (right) OHCs to the ramp-like deflections of the stereocilia bundle with a fluid-jet (bottom) at different holding potentials from -90 to +110 mV. For clarity, MET current traces at different potentials are equally spaced and the voltage-gated conductance is subtracted. Inset: Bright-field image of the organ of Corti explant with the fluid-jet deflecting OHC bundle (bottom) and a recording patch clamp pipette underneath the focal plane (right). (B) Voltage-dependence of the normalized MET currents in control and *Cib2^R186W/R186W^* OHCs. Mean±SE are shown. (C) Normalized MET current – hair bundle displacement relationships (I-X curves) at negative (-90 mV) and positive (+110 mV) potentials for the cells shown in (A). Data were fit to a double Boltzmann curve. (D) Resting open MET channel probability at different holding potentials (-90 mV to +110 mV). Asterisks indicate significance: **, P<0.01; ***, P<0.001 (two-way ANOVA). Number of cells/mice: control, n=16/6 (*Cib2^+/+^*, n=12/4; *Cib2^+/R186W^*, n=4/2); *Cib2^R186W/R186W^*, n=9/2. (G) Additive effects of R186W variant and cell depolarization on the leftward shifts of the I-X curves. Each I-X curve is represented as a data point with the X-coordinate determined at 20% of MET channel activation. Black and red points correspond to the data at holding potential of -90 mV for control and *Cib2^R186W/R186W^*OHCs, correspondingly. Grey and light red points correspond to the same OHCs at +110 mV holding potential. Age of the cells: P4-P6, all of them were located approximately in the middle of the apical turn of the cochlea.

Consistent with the rigid probe data, our fluid-jet experiments also showed a decrease in the width of I-X curves in *Cib2^R186W/R186W^* OHCs compared to the control **(Figure 3E, Supplemental Fig. 3B)**. In contrast to the rigid probe, fluid-jet is effective in deflecting hair bundles in both positive and negative directions. Therefore, we were also able to quantify the open MET channel probability at resting bundle position (P_open_) and found that *Cib2^R186W/R186W^* OHCs had significantly larger P_open_ compared to the control OHCs **(Figure 3F, Supplemental Fig. 3C)**. These two results, a steeper I-X curve and an increased Popen, could be easily reconciled by a model that assumes that R186W variant of CIB2 makes the MET channel more sensitive to the external force.

### AlphaFold2 modeling provides an explanation of how the R186W variant may increase the sensitivity of MET channels to the external force

A structural model of CIB2-TMC1/2 interactions was generated using Alphafold multimer (**Figure 3G**). The heterotetrameric complex was predicted with high confidence (**Supplemental Figure 4**), and observed interfaces were highly similar to the recently determined structure of the TMC1/CALM-1 complex from C. elegans ^34^. Analysis of the molecular interfaces in the complex reveal that one copy of CIB2 integrates into the complex in tight association with TMC1 while another copy of CIB2 integrates with TMC2. In both cases, the interfaces are highly similar and tightly integrated with half of CIB2 surface residues buried at the interface and extensive interactions. Analysis reveals 22 hydrogen bonds and 21 salt bridges for CIB2-TMC1 and 24 hydrogen bonds and 23 salt bridges for CIB2-TMC2. The extensive interaction interface between CIB2 and TMC1/2 may confer mechanical constraints to the TMC proteins, counteracting the external force and setting the “optimal” sensitivity of the MET channel. Of particular interest, in the CIB2-TMC1 interface R186 forms a salt bridge with TMC1 E172 (**Figure 3H, left**). In the CIB2-TMC2 interface, R186 forms a salt bridge with TMC2 E224 (**Figure 3H, right**). This is consistent with the hypothesis that the p.R186W mutation releases some mechanical constraints imposed by CIB2 to the MET channel, increasing P_open_ and the force sensitivity (**Figure 3I**).

### Exaggerated voltage-dependence of MET currents in *Cib2^R186W/R186W^*OHCs

It is well known that depolarization of a hair cell results in the leftward shift of I-X curve and the increase of P_open_ that are thought to be caused by the decrease of Ca^2+^influx into the cell through the MET channels ^35, 36^. Since CIB2 is a potential “Ca^2+^ sensor” within MET complex, we explored the effects of R186W variant of voltage dependence of the MET currents. We recorded MET currents evoked by identical fluid-jet ramp stimuli but at different holding potentials (from -90 to +110 mV) **(Figure 4A)**. With Cs-based intracellular solution, *Cib2^R186W/R186W^* OHCs exhibit MET currents with more prominent rectification compared to the control OHCs, while reversal potential was similar (**Figure 4B**). As expected, in the control OHCs, we observed a leftward shift of the MET current-displacement curves induced by cell depolarization **(Figure 4C, left)**. Interestingly, I-X curves already shifted to the left in *Cib2^R186W/R186W^* OHCs at negative holding potentials shifted even further at positive potentials **(Figure 4C, right)**. As a result of this exaggerated shift, the difference in P_open_ between control and *Cib2^R186W/R186W^* OHCs became even larger at positive potentials **(Figure 4D)**. The effects of R186W variant and cell depolarization were additive *i.e.*, R186W produced a relatively small leftward shift of I-X curves compared to the control at negative potentials, while cell depolarization produced further exaggerated leftward shift in *Cib2^R186W/R186W^* OHCs **(Figure 4E)**. Thus, R186W variant of CIB2 does not disrupt the voltage-dependent machinery within the MET complex but modulates its function.

### R186W variant slows down MET channel activation and disrupts fast adaptation

To explore the effects of R186W variant on the dynamics of the MET currents, we performed patch-clamp experiments using a microfabricated rigid probe capable to produce high-speed bundle deflections of ∼1 μm in less than 10 µs **(Figure 5A)**. The use of this microfabricated probe allow us to resolve the two-state kinetics of MET channel activation in wild type OHCs, which was not possible with slower “conventional” probe **(Figure 5B)**. The speed of the microfabricated probe movement was significantly faster than the rising phase of the MET currents in the wild type OHCs (**Figure 5D**). Interestingly, the time constant of MET channel activation in *Cib2^R186W/R186W^*OHCs was significantly larger than that in wild type OHCs **(Figure 5C,D)** at all tested ages of OHCs (**Supplemental Fig. 3D**). This difference in time constants of MET channel activation also did not depend on the amplitude of the MET current – even the smallest MET responses to small bundle deflections activated faster in wild type OHCs compared to *Cib2^R186W/R186W^*OHCs (**Supplemental Fig. 5A**). We concluded that R186W variant slows down MET channel activation either directly or indirectly.

**Figure 5.**
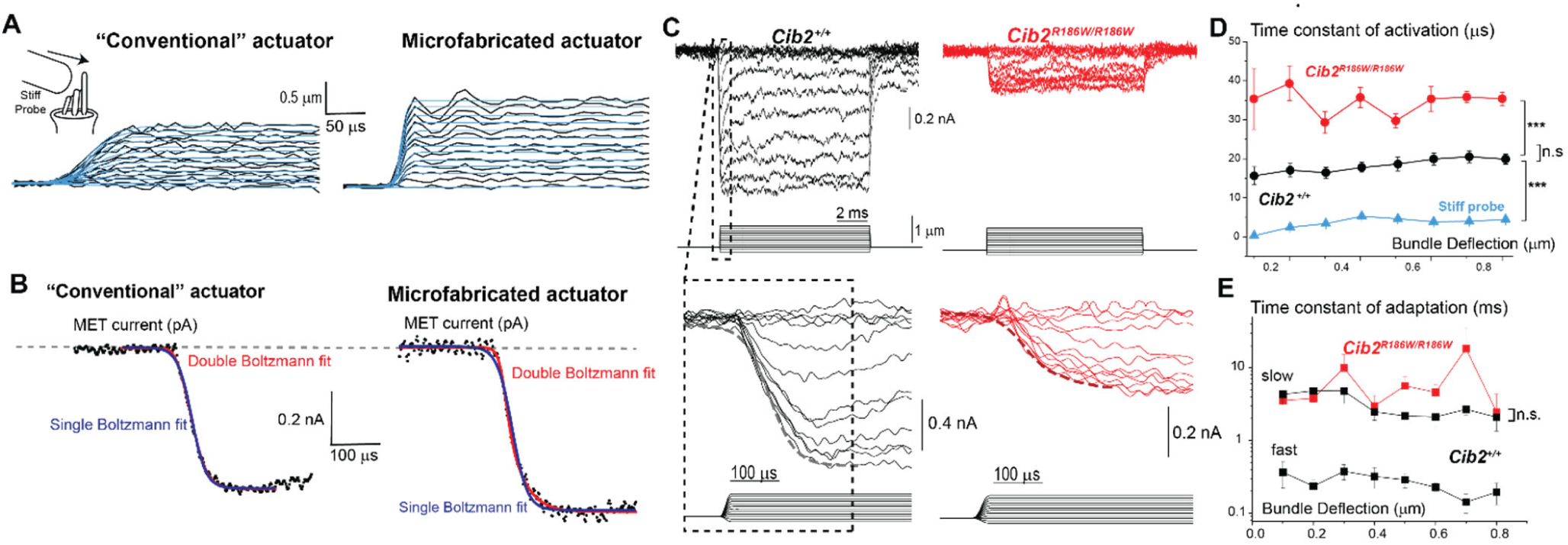
R186W variant of CIB2 slows down MET channel activation and eliminates fast adaptation. (A) Step movements of a conventional (left) and a microfabricated (right) piezo-driven stiff-probes obtained from frame-by-frame analysis of video recordings at 90,000 frames per second. Time constants of these probe movements (blue lines) obtained with a single Boltzmann fit (see methods) are: τ = 15.34±1 µs and τ = 4.40 ± 0.76 µs, correspondingly. (B) Only MET currents evoked by a faster microfabricated probe (right) but not a conventional (left) probe revealed two-stage activation of MET channels. Initial phase of MET responses to the saturated stereocilia bundle deflections were fit to single (blue line) or double Boltzmann (red line) functions. Results from Alaikés Bayesan and F-test confirmed that only the currents elicited by the microfabricated probe are better fit with the double-Boltzmann function. (C) Representative traces of the MET currents evoked by step-like deflections of the hair bundle using the microfabricated probe in wild type (left) and *Cib2^R186W/R186W^* (right) OHCs (top panel) on slower (top) and faster (bottom) time scales. (D) Average time constants of activation of the MET current in wild type (black) and *Cib2^R186W/R186W^* (red) OHCs at different bundle displacements. Due to the slow MET channels activation in *Cib2^R186W/R186W^* OHCs the fit to a single Boltzmann function was used in both genotype groups. Boltzmann function matches the step response of the Bessel filter used to filter the command voltage driving the piezo-driven probe (see Methods). Number of cells/mice: *Cib2^+/+^*, n=16/12; *Cib2^R186W/R186W^*, n=6/6. (E) Time constants of adaptation were obtained by fitting the MET current decay with a double (wild type, fast and slow adaptation) or a single (*Cib2^R186W/R186W^*) exponential function. Number of cells/mice: *Cib2^+/+^*, n=7/4; *Cib2^R186W/R186W^*, n=11/6. (D,E) Data are shown as mean±SE. Asterisks indicate statistical significance of the differences: ***, P<0.001; n.s., non-significant (two-way ANOVA). Age of the cells: P4-P6, all of them were located approximately in the middle of the apical turn of the cochlea.

Despite the small amplitude of MET currents, *Cib2^R186W/R186W^*OHCs showed some adaptation **(Figure 4C and Supplemental Fig. 5B)**. However, unlike wild type OHCs, in the vast majority of *Cib2^R186W/R186W^*OHCs, it was not possible to use the double exponential fit to quantify fast and slow adaptation (**Supplemental Fig. 5C**). Therefore, a single exponential function was used instead in *Cib2^R186W/R186W^*OHCs. Interestingly, the time constant of slow adaptation in wild type OHCs was almost identical to the time constant of adaptation in *Cib2^R186W/R186W^* OHCs determined by a single exponential fit **(Figure 5E)**. We concluded that R186W variant disrupts fast adaptation.

We also explored potential voltage-dependent changes of the time constant of MET channel activation in wild type and *Cib2^R186W/R186W^*OHCs. MET currents were evoked by identical step-like bundle deflections at negative (-90 mV) and positive (+90 mV) holding potentials in the same cells **(Figure 6A)**, while the time constant of MET activation was determined from a fit to a single Boltzmann function. We observed a trend toward slower MET activation at positive holding potentials in both wild type and *Cib2^R186W/R186W^*OHCs **(Figure 6B)**. However, this trend was not statistically significant at P=0.01 level and, therefore, we cannot yet draw any meaningful conclusions.

**Figure 6:**
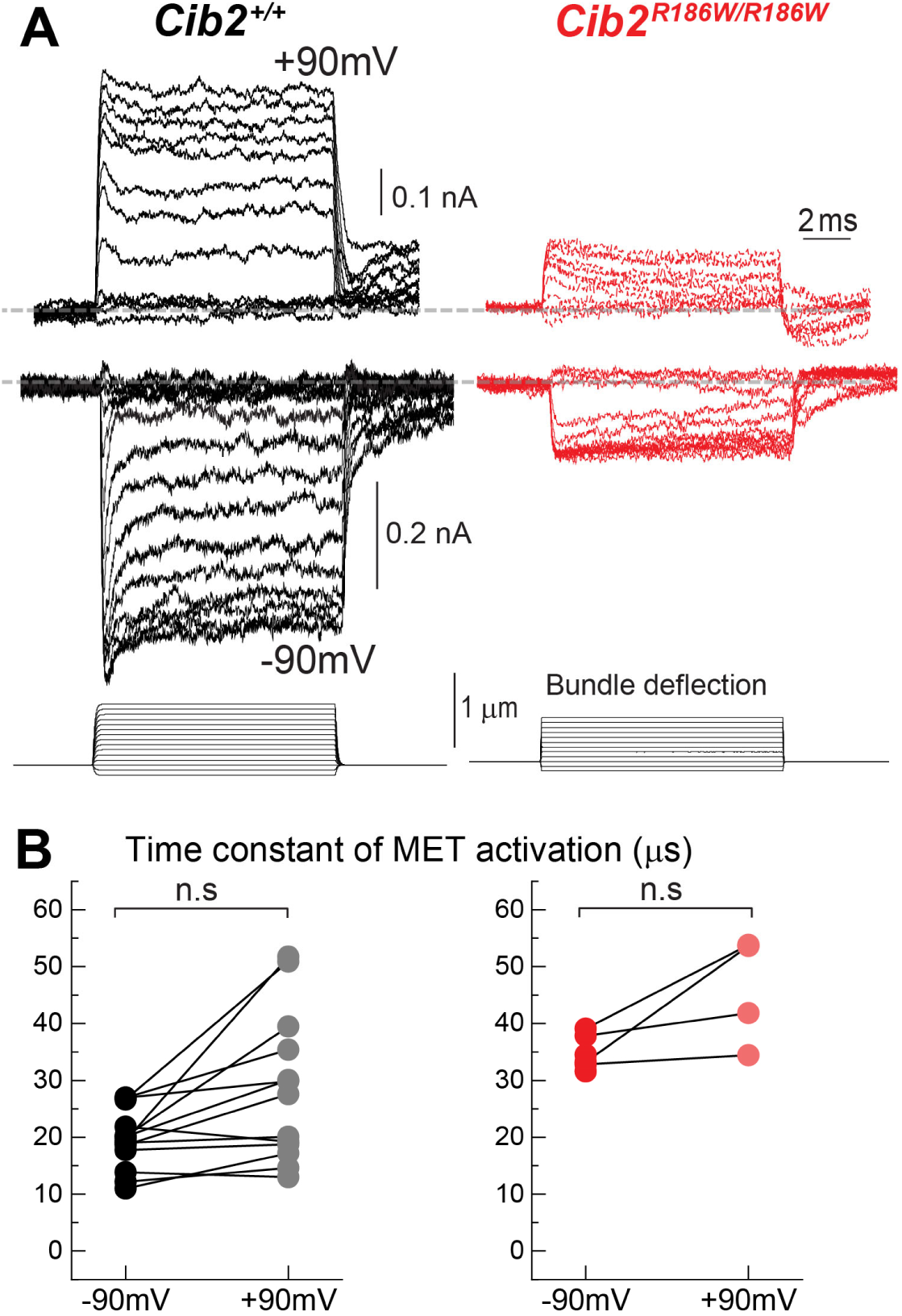
Non-significant trend toward depolarization-induced increase of the time constant of MET channel activation in both *Cib2^+/+^* and *Cib2^R186W/R186W^* OHCs. (A) Representative MET currents evoked by step-like deflections of the stereocilia bundle with the stiff probe in the same *Cib2^+/+^* (left) and *Cib2^R186W/R186W^* (right) OHCs at two holding potentials: +90 mV (top) and -90 mV (bottom). (B) Time constants of MET channel activation at positive and negative holding potentials in the same *Cib2^+/+^* (left) and *Cib2^R186W/R186W^* (right) OHCs. Each data point represents one cell and was calculated as an average of the time constants at bundle deflections of 0.1-0.8 µm (see Fig. 5D). Statistics: n.s, p>0.01 (paired samples *t*-test). Number of cells/mice: *Cib2^+/+^*, n=13/11; *Cib2^R186W/R186W^*, n=4/4

### R186W variant disrupts lower tip link density and mis-localizes BAIAP2L2 from stereocilia tips

Slower activation of MET channels in *Cib2^R186W/R186W^*OHCs is hard to understand if the only effect of p.R186W mutation is the increased sensitivity of the channel to the external force (**Figure 3I**). Therefore, we explored potential ultrastructural changes at the tips of hair cell stereocilia using Focus Ion Beam-Scanning Electron Microscopy (FIB-SEM). The 20 nm serial sectioning allows us to visualize the lower tip link density in practically every second row stereocilium in the wild type IHCs (**Figure 7A, left**). This electron-dense structure presumably harbors the components of the MET machinery. In the *Cib2^R186W/R186W^* IHCs, however, the lower tip link density was no longer visible as a cohesive structure. Instead, the electron-dense material was dispersed throughout the entire region at the tip of stereocilium (**Figure 7A, right**).

**Figure 7:**
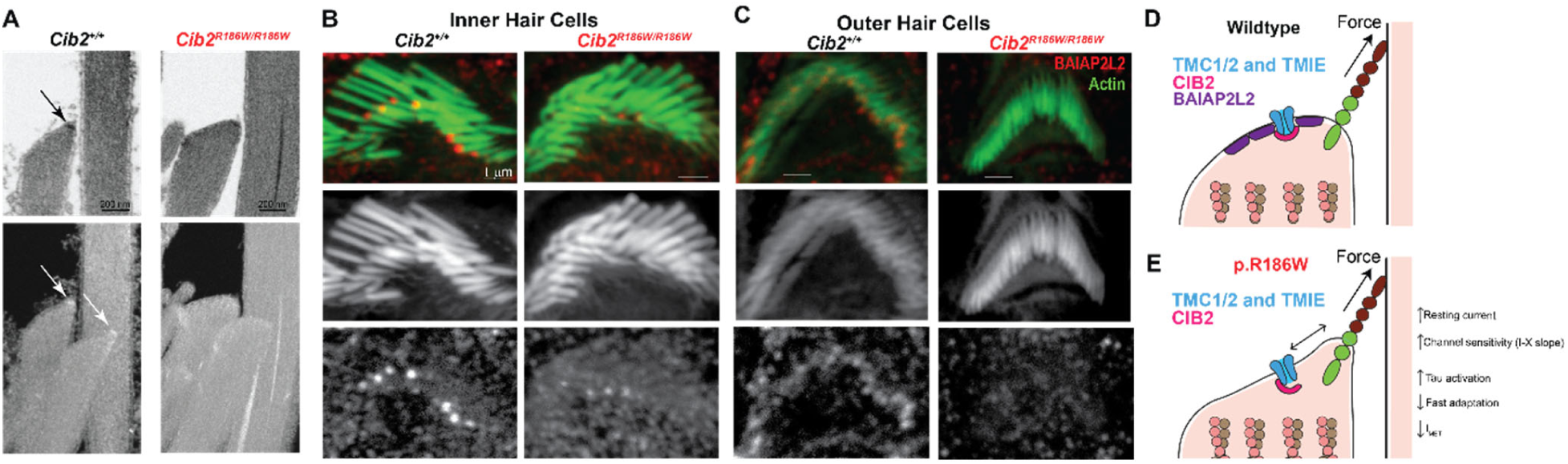
R186W variant causes disruption of “lower tip link density” and mis-localization of BAIAP2L2. (A) Individual Focused Ion Beam-Scanning Electron Microscopy (FIB-SEM) sections (top) and maximum intensity projections of several FIB-SEM sections encompassing two transducing 2^nd^ row stereocilia (bottom) in wild types (left) and *Cib2^R186W/R186W^* (right) IHCs. Arrows point to the “lower tip link densities” that are not apparent in *Cib2^R186W/R186W^* IHCs anymore. Age of the cells: P16. (B-C) Maximum intensity projections of BAIAP2L2 immunolabeling in IHCs (B) and OHCs (C) of *Cib2^+/+^* (left) and *Cib2^R186W/R186W^* (right) mice. The top panels show merged images of BAIAP2L2 (red) and F-actin (green). Middle and bottom panels show individual channels: F-actin (middle) and BAIAP2L2 (bottom). Age of the cells: P7. (D) BAIAP2L2-dependent reinforcement of the plasma membrane may be essential for the proper force transmission to the MET channels. (E) The loss of BAIAP2L2 from the tips of stereocilia in *Cib2^R186W/R186W^* mice may decrease the efficiency of force transmission to the MET channels, resulting in their slower activation.

An attractive candidate for shaping the tips of stereocilia is BAR/IMD Domain Containing Adaptor Protein 2 Like 2 (BAIAP2L2). BAR domain proteins typically bind to the plasma membrane, thereby reinforcing the membrane and contributing to its curvature and mechanical stiffness ^37^. BAIAP2L2 is localized to the tips of transducing stereocilia and essential for their maintenance ^38, 39^. Furthermore, it has been reported that BAIAP2L2 interacts with CIB2 and is mis-localized in *Cib2* knockout mice ^39^. Typically, BAR domain proteins self-polymerize and form large membrane-associated complexes ^37^, which may be essential for formation of the lower tip link density in stereocilia. Therefore, we examined the potential effect of p.R186W mutation in CIB2 on BAIAP2L2 localization in the auditory hair cell stereocilia using immunofluorescent labeling and confocal imaging in P7 cochleae of *Cib2^R186W/R186W^*and *Cib2^+/+^* mice. In wild type IHCs and OHCs, BAIAP2L2 was concentrated at the tips of transducing shorter row stereocilia and within the cuticular plate (**Figure 7B,C, left**). However, BAIAP2L2 was not present anymore at the stereocilia tips in *Cib2^R186W/R1^*^86^ hair cells (**Figure 7B,C, left**). Thus, we hypothesized that BAIAP2L2-dependent structures are essential for the proper force transmission to the MET channel (**Figure 7D**). Mis-localization of BAIAP2L2 from the tips of transducing stereocilia in *Cib2^R186W/R186W^*OHCs would disrupt these structures, resulting in the force transmission largely through viscoelastic plasma membrane, which would result in slower opening of the MET channel (**Figure 7E**).

## DISCUSSION

To the best of our knowledge, this is the first study that systematically explores the time constant of MET activation in mammalian auditory hair cells. A significant decrease of this time constant was observed in *Cib2^R186W/R186W^* OHCs, independent of the amplitude of the MET current and the age of the animals. This decrease may be caused either by the slower gating of MET channel itself or by the slower force transmission to the channel. Irrespective of the exact molecular composition of the MET channels studied here (see discussion below), slower gating of the MET channels (Figure 5D) is hard to reconcile with their increased sensitivity to hair bundle deflection (Figure 3C-E). On the other hand, we found that the effects of p.R186W mutation in CIB2 are not limited to the MET channels but also involve disruption of stereocilia tip density (presumably containing the MET complex), loss of BAIAP2L2 from the tips of transducing stereocilia (Figure 7), and over-elongation (or “tenting”) of the second row stereocilia (Figure 2B,D). One may argue that all these effects may be secondary to the long-term decrease in MET current. Indeed, while the effects of long-term MET channel blockage on stereocilia tip density have not yet been investigated, the loss of BAIAP2L2 has been already reported ^40^. However, the overgrowth of 2^nd^ row stereocilia in *Cib2* mutants (Figure 2E, see also ^18^) cannot be explained by decreased MET current since blockage of the MET channels normally causes retraction of these stereocilia but not their growth ^41^. Thus, CIB2 seems to have a role in shaping stereocilia tips that is independent of its effects on MET channels. It was also reported that CIB2 directly binds BAIAP2L2, and BAIAP2L2 is mis-localized from the tips of stereocilia in *Cib2* knockout mice ^39^. However, CIB2 is present in the stereocilia of *Baiap2l2* knockout mice ^39^, while young postnatal auditory hair cells of these mice have apparently normal amplitude of MET currents ^38^. Thus, it is more likely that CIB2 controls BAIAP2L2 localization, thereby controlling the mechanical properties of the plasma membrane in the vicinity of the MET channels and the effectiveness of force transmission to these channels.

As a potentially Ca^2+^-sensitive element, CIB2 may participate in the dynamic regulation of mechanical force within the MET machinery known as adaptation (reviewed in ^42, 43^). Adaptation has been historically separated into myosin-dependent “slow” adaptation ^44–46^ and a “fast” component, the mechanisms of which are still obscure ^2, 47^. One of the models, a “tension-release" model of fast adaptation, postulates that a Ca^2+^-dependent element that is mechanically in series with the MET channel undergoes a conformational change upon Ca^2+^ binding, thereby decreasing the tension within MET machinery ^48, 49^. While there is limited evidence for the existence of a tension-release element in vestibular hair cells ^50^, its existence and molecular identity in the mammalian auditory hair cells has never been determined. In theory, CIB2 may function as such element and, on the first sight, the loss of fast adaptation in *Cib2^R186W/R186W^* OHCs (Figure 5E) confirms this hypothesis. However, we would refrain yet from making such a conclusion, first, because slowing of MET channel activation (Figure 5C-D) would certainly mask fast adaptation and, second, because fast adaptation in mammalian auditory hair cells may not depend on Ca^2+^ ^51^.

It is not easy to decipher the effects of R186W variant of CIB2 on the MET channels themselves. After we collected most of the data of this study, a new report has emerged, which demonstrated that CIB2 is essential for proper trafficking of TMCs to the tips of stereocilia ^19^. Based on an observation that R186W variant affects stereocilia localization of TMC1 more than that of TMC2 and on disappearance of MET responses in the double *Tmc2^-/-^;Cib2^R186W/R186W^* mutants, the authors proposed that MET responses in *Cib2^R186W/R186W^* hair cells are mediated by the channels containing only TMC2. If this is the case, one would expect very similar MET responses in *Cib2^R186W/R186W^* OHCs on a wildtype background (expressing both TMC1 and TMC1) and on *Tmc1^dn/dn^* background (without TMC1). If not, the differences between MET responses in *Tmc1^+/+^*;*Cib2^R186W/R186W^* and *Tmc1^dn/dn^*;*Cib2^R186W/R186W^* OHCs may reveal the effects of R186W variant on TMC1-containing MET channels, which is something that cannot be accessed directly since *Tmc2^-/-^;Cib2^R186W/R186W^*OHCs have no measurable MET currents ^19^. Obviously, immunofluorescent imaging of tagged TMC proteins utilized in ^19^ may have limited ability to detect just a few TMC1-containing channels that are expected to remain in Cib*2^R186W/R186W^* OHCs on a wildtype background.

In this study, we have analyzed developmental changes of single MET channel conductance and limited our analysis to the apical OHCs of wildtype mice at P4-P6, right before single MET channel conductance start to increase at P7 (Supplemental Figure 3A), presumably due to an increased number of MET channels containing TMC1 subunits only ^52^. Indeed, it is known that the apical OHCs at P4-P6 express both TMC1 and TMC2 ^52^. Therefore, we could compare our data with a recent study that also explored MET currents in presumably apical OHCs but in *Tmc1^dn/dn^*;*Cib2^R186W/R186W^* mice expressing TMC2 only ^19^. A common finding between these two studies is the increase of the open probability of MET channels at rest (P_open_) in OHCs expressing R186W variant of CIB2. This increase remained unexplained in the previous study but was linked to the steeper I-X curves in our study (Figure 3C-I). Besides changes in P_open_, MET currents in *Tmc1^dn/dn^*;*Cib2^R186W/R186W^*were apparently normal with a prominent fast adaptation and practically unchanged dependence of P_open_ on the intracellular potential ^19^. In contrast, *Cib2^R186W/R186W^* OHCs in our study exhibited no measurable fast adaptation and exaggerated voltage-dependence of P_open_ (Figure 5C,E; Figure 4D-E). Although some unanticipated differences between two independently generated *Cib2^R186W^* strains of mice cannot be excluded (*e.g.*, differences in genetic background that may influence mutated MET channels ^53^), the most plausible explanation is that our study reveals CIB2 effects on more mature heteromeric TMC1/TMC2 channels. This explanation is also consistent with the results of AlphaFold2 modeling that predicts nearly identical interaction interfaces of CIB2 with TMC1 and TMC2 (Figure 3G-H), suggesting again that the previously unreported effects of R186W variant on MET currents observed in our study are likely to be due to the presence of TMC1 subunits.

Another striking effect of R186W variant on MET currents is the increase of the slope of I-X curves that was observed with both rigid probe and fluid-jet deflections of the hair bundles (Figur. 3D-E). One could argue that the slope of I-X curve depends on OHC bundle morphology that is changing in *Cib2^R186W/R186W^* mice (Figure 2). However, the loss of staircase architecture of the hair bundle and the larger variability of the heights of transducing stereocilia should make the I-X curve shallower rather than steeper ^32^. Likewise, if anything, the overgrowth of transducing stereocilia in *Cib2^R186W/R186W^* OHCs (Figure 2C,E) would make tip links less inclined, which in turn would result in shallower rather than steeper I-X curve ^54^. On the other hand, the AlphaFold2-based model postulating that R186W variant of CIB2 releases some (but not all) mechanical constrains on TMC1/2 conformations induced by an external force (Figure 3I) provides a mutually consistent explanation for both the steeper I-X curve and increased P_open_ in *Cib2^R186W/R186W^* OHCs.

Because CIB2 is the only potentially Ca^2+^-dependent MET channel component known so far, it is an attractive candidate for mediating at least some of the multiple effects of Ca^2+^ on hair cell mechanotransduction. It has been argued that, at the normal intracellular concentrations of free Ca^2+^ and Mg^2+^, EF-hand domains of CIB2 should be occupied by Mg^2+^ rather than by Ca^2+^ ^19^. However, this argument may not work in a hair cell since concentration of Ca^2+^ at the tips of transducing stereocilia may be significantly higher in physiological conditions ^55^. On the first sight, our data also suggest that CIB2 is not a “Ca^2+^ sensor” of the MET complex, because p.R186W mutation in CIB2 does not eliminate the dependence of P_open_ on cell depolarization (Figure 4), which is generally assumed to be a result of the decreased Ca^2+^ influx through the MET channels ^35, 36^. However, this dependence was exaggerated in *Cib2^R186W/R186W^* OHCs indicating that R186W variant does have a modulatory effect. It can be explained if CIB2 has indeed multiple interaction sites with TMC1 and TMC2 as predicted by AlphaFold2 modeling (Figure 3G; Supplemental Figure 4) and they remain stable at different Ca^2+^ concentrations ^19^. Then, Ca^2+^-dependent conformational changes in CIB2 may exert forces to TMC1/2, thereby changing the open channel probability at rest. Disruption of CIB2-TMC1/2 interaction outside EF-hand domains (*e.g.*, at R186 site) wouldn’t affect Ca^2+^ binding to CIB2 but it may remove mechanical constrains to TMC1/2 and exaggerate intramolecular movements of TMC1/2 forced by Ca^2+^-dependent CIB2 conformations. Obviously, additional structural data are needed to test these speculations.

## ACKNOWLEDGEMENTS

This study was supported by NIDCD/NIH R01DC012564 (to Z.M.A. and G.I.F.) and by OD/NIH S10OD025130 (to G.I.F). Electron microscopy was performed at the Electron Microscopy Center, which belongs to the National Science Foundation NNCI Kentucky Multiscale Manufacturing and Nano Integration Node, supported by ECCS-1542174. We thank Dr. Catalina Velez-Ortega (University of Kentucky) for the access to Leica SP8 confocal microscope, and Drs. Julia Halford and Peter Barr-Gillespie (Oregon Health & Science University) for their advice with BAIAP2L2 immunolabeling.

## METHODS

### Animals

*Cib2^R186W^* mice: *Cib2^R186W^* mouse model carrying the DFNB48 p.Arg186Trp (R186W) pathogenic variant was generated using CRISPR/Cas9 technology. Mice carrying the targeted allele were maintained on C57Bl6/J background and crossed with wild type mice to get rid of off-target variations. *Cib2^R186W^* allele was then confirmed by sequencing. All animal procedures were approved by the Institutional Animal Care and Use Committees (IACUCs) at University of Maryland (protocol #0420002) and University of Kentucky (protocols 00903M2005 and 2019-3414).

### Patch clamp recordings of the MET currents

Organ of Corti explants were dissected at postnatal day 4 through 7 (P4-P7) in Leibovitz’s L-15 cell culture medium (Invitrogen), containing the following inorganic salts (in mM): NaCl (137), KCl (5.4), CaCl_2_ (1.26), MgCl_2_ (1.0), Na_2_HPO_4_ (1.0), KH_2_PO_4_ (0.44), MgSO_4_ (0.81). Then, they were mounted on a custom-made chamber that has a thick glass bottom plate, single stranded dental floss filaments for securing explants in place, and low-profile walls for shallow electrode access to the sample. The hair cells were observed either with Nikon E600 or Olympus BX51WI upright microscopes, equipped with either 100X, 1.1 NA or 100X, 1.0 NA long working distance water-immersion objectives, correspondingly. The samples were continuously perfused with fresh L-15 solution via a pipette with ∼50-100 μm diameter located in the vicinity of the field of view. Patch pipettes were made of thick-wall borosilicate capillaries and had resistances of 3-6 MΩ in the bath. We used two Cs-based intra-pipette solutions contained (in mM): i) CsCl (147), MgCl_2_ (2.5), EGTA (1.0), K_2_ATP (2.5), and HEPES (5) in the experiments with fluid-jet bundle deflection; and ii) CsCl (114), MgCl_2_ (3.5), EGTA (1.0), Na_2_ATP (5.0), HEPES (10), creatine phosphate (5.0), and ascorbic acid (2.0) in the experiments with rigid probe deflection of the hair bundles. Series resistance was compensated (up to 60%) only in the experiments with rigid probe stimulation, because relatively slow hair bundle deflections with a fluid-jet (∼1 ms rise time) resulted in a rather slow rise of MET current, which did not require a fast voltage clamp. However, the voltage drop across series resistance was subtracted in all experiments to obtain correct intracellular potentials. Patch clamp recordings were performed with a MultiClamp 700B amplifier and pClamp 11 software (Molecular Devices, Sunnyvale, CA). Hair cells were held at a potential of -60 mV between the recordings and at -90 mV for the transient periods of MET recordings, unless stated otherwise. All recorded hair cells were located approximately in the middle of the apical turn of the cochlea.

### Hair bundle deflection with fluid-jet

Pressure was generated using a High Speed Pressure Clamp (HSPC-1, ALA Scientific) and applied to the back of a ∼5 μm thin wall pipette filled with the L-15 bath solution. The pipette tip was positioned at ∼8 μm in front of the hair bundle. It was determined that the force generated by this microjet depends linearly on the applied pressure ^56^. Before each experiment, the steady-state pressure was adjusted to zero by monitoring debris movement in front of a fluid-jet. Hair bundle movement was recorded with CS2100M-USB sCMOS camera (ThorLabs, Newton, New Jersey) at a rate of 166-270 frames per second, triggered by pClamp. The “frame ready” signal from the camera was recorded by pClamp for synchronization of the current and video records. Off-line frame-by-frame computation of hair bundle displacements was performed using MATLAB scripts developed for quantifying electromotility of isolated OHCs ^57^. Briefly, the intensity profiles across the middle of the bundle were collected for each frame of the video record. The number of points in these profiles were interpolated 50 times to obtain sub-pixel resolution. The frame-by-frame shifts of the intensity profiles, and hence bundle position, were determined by the least square algorithm. The typical error of the movement measurements with this technique is ∼20 nm. The slow drift of the bundle position, if present, was corrected. At this point, a hair bundle displacement – fluid-jet pressure ΔX(P) relationship was determined for each recording and was used for constructing MET current – bundle displacement (e.g. “I-X”) curves. The slope of ΔX(P) relationship at the small bundle deflections is proportional to mechanical compliance of the hair bundle, while reciprocal of this value is proportional to the bundle stiffness.

### Hair bundle deflection with a stiff probe

OHC bundles were also deflected with a borosilicate-glass pipette probe that was driven by a calibrated piezoelectric stack actuator (PL055.31, Physik Instrumente, Germany). The pipette was heat-polished to a diameter of ∼5-7 µm, matching the shape of the OHC bundle, which ensures simultaneous deflection of the maximum number of stereocilia. The protruding part of the probe was 3-5 mm long, which alleviated lateral resonances. The stimuli for hair bundle deflections were generated by pClamp, low-pass filtered at 50 kHz with a Bessel-pole filter, and then amplified by the piezo-amplifier (PX200, PiezoDrive, Callaghan, Australia) that drives the piezo-actuator. The speed of the microfabricated probe was tested by capturing videos of the step-like movements from 0 to 1 µm, by 0.2 µm increments at 90.000 fps using a VEO-E 310L high speed camera (Phantom/Ametek). Movement of the probe was quantified by off-line frame-by-frame analysis, the same way as we did for the quantification of hair bundle movements described above. The speed of the new microfabricated probe was compared with a “conventional probe” driven by a PK4FA2H3P2 (ThorLabs) piezo stack.

### MET current analysis

Using OriginPro 2020 software (OriginLab, Northampton, MA), I-X curves obtained either from step or ramp deflections of the hair bundles were fit to the double Boltzman function: I_MET_ = I_MAX_/(1+exp(A1×(P1-ΔX))×(1+exp(A2×(P2-ΔX))))-I_MIN_, where ΔX is the bundle displacement, I_MAX_ is the maximal MET current, I_MIN_ is the whole cell current at saturating negative bundle deflection when all MET channels are closed, while A1, A2, and P1, P2 are the parameters determining the slope and positioning of the I-X curve.

Single MET-channel properties were determined from nonstationary current noise^58^. The hair bundle was deflected from a saturating negative to a saturating positive position by several consecutive and identical slow ramps of 250 ms duration. The resulting MET current traces were digitized at 10 kHz and low-pass filtered at 1.6 kHz to reduce noise and reveal trace-by-trace variations. If the set of records showed unstable MET current at saturating positive or negative displacements, it was discarded. Then, the relationship between MET current variance σ (corrected by subtracting the variance attributable to the background noise) and the mean MET current (I) was fit with the parabolic equation: σ(I) = I_0_×I − I^2^/N_MET_, where I_0_ is the single-channel current and N_MET_ is the number of channels per bundle ^58^.

The data from the stiff probe were obtained by 0.2 µm step-like deflection of the stereocilia bundle (from -0.2 to 1 µm). MET currents were recorded using a 30 kHz low pass filter. To determine the time constant of activation, only the first millisecond of the current recording was used to fit the onset of the current with a single-Boltzmann function. For fast and slow adaptation, a double exponential fit was used to follow the current decay in the first 5-7 ms of the current response.

### CIB2 immunostaining

The cochlear sensory epithelia were isolated, fine dissected and permeabilized in 0.25% Triton X-100 for 1 h and blocked with 10% normal goat serum in PBS for 1 h. The tissue samples were probed with primary antibody overnight and after three washes were incubated with the secondary antibody for 45 min at room temperature. Rhodamine phalloidin or Alexa fluor phalloidin 488 were used at a 1:250 dilution for F-actin labeling. Nuclei were stained with DAPI (Molecular Probes).

### BAIAP2L2 immunolabeling

BAIAP2L2 immunolabeling followed the previously published protocol ^40^. Briefly, the temporal bones were dissected, and the cochleae were exposed. Small holes were made at the base and apex of the cochlea to allow perfusion of the fixative. The cochleae were then perfused with 4% PFA and fixed for 25 minutes. Two washes of 10 minutes in PBS were done before dissecting the sensory epithelia. The tissue was permeabilized using 0.2% Triton X-100 in PBS for 10 minutes at room temperature. The tissues were transferred to a blocking solution (5% horse serum in PBS) for one hour at room temperature. Then, the organ of Corti explants were incubated overnight with the primary antibody: anti-BAIAP2L2 (Sigma-Aldrich Cat# HPA003043,RRID:AB_2227864) diluted 1:150 in the blocking solution. Two washes of 10 minutes were done before incubating the samples for four hours at room temperature with the secondary antibody and Phalloidin-Alexa Fluor 488 counterstain. The secondary donkey anti-rabbit 568nm (2 mg/ml) antibodies were diluted 1:1000 with blocking solution, while Phalloidin was diluted 1:1000 with the same solution. After the incubation, three more washes of 10 minutes in PBS were done before mounting the samples.

Images were acquired using either a LSM 700 laser scanning confocal microscope (Zeiss, Germany) with a 63X 1.4 NA or 100X 1.4 NA oil immersion objectives or Leica SP8 laser scanning confocal microscope with a 100X 1.44 NA objective lens. Stacks of confocal images were acquired with a Z step of 0.05-0.5 µm and processed using ImageJ software (National Institutes of Health). Experiments were repeated at least 3 times, using at least three different animals.

### Scanning Electron Microscopy (SEM) and tip link quantification

*Cib2^R186W^* mutant cochleae were fixed in 2.5% glutaraldehyde in 0.1 M cacodylate buffer, pH 7.4 (Electron Microscopy Sciences, Hatfield, PA) supplemented with 2 mM CaCl_2_ (Sigma-Aldrich) for 1–2 h at room temperature. Then, the sensory epithelia were dissected in distilled water, dehydrated through a graded series of ethanol, critical point dried from liquid CO_2_ (Leica EM CPD300), sputter-coated with 5 nm platinum (Q150T, Quorum Technologies, Guelph, Canada), and imaged with a field-emission scanning electron microscope (Helios Nanolab 660, FEI, Hillsboro, OR). Tip link count was performed in the samples that were used for patch clamp recordings and then fixed at the end of the experiment. A “tip” link was defined as a link that extends obliquely from the top of a lower row stereocilium to the side of a taller stereocilium in the direction of mechanosensitivity of the bundle. Any other link originating at the hemisphere of the tip of a shorter stereocilium was not considered as a tip link. See ^59^ for more details.

### Serial sectioning Electron with Focused Ion Beam and Scanning Electron **Microscopy (FIB-SEM)**

Cochleae were extracted from temporal bones and gently perfused through the oval window with a solution containing 2.5% glutaraldehyde, 2% paraformaldehyde in 0.1M cacodylate buffer, pH=7.4 (Electron Microscopy Science, cat# 15960-01), and supplemented with 2mM CaCl_2_ and 1% Tannic acid (Electron Microscopy Sciences, cat# 21710). The cochleae were kept in this fixative overnight at 4°C, and then the fixative was diluted ∼1:5 with 0.1M cacodylate buffer. Then, the organs of Corti were dissected in distilled water with the tectorial membrane kept intact and high-pressure frozen with Leica EM ICE high-pressure freezer. The frozen samples were transferred to Leica EM AFS2 freeze substitution machine and kept in methanol with 1% uranyl acetate at -90°C for 33 hours and then slowly warmed at about 4°C per hour to -45°C. Samples were then washed with fresh 100% methanol for 24 hours at -45°C to fully replace uranyl acetate. Then the methanol was gradually replaced with Lowicryl HM-20 resin (monostep embedding kit, Electron Microscopy Sciences, cat# 14345): 50% Lowicryl (2 hours), 75% Lowicryl (overnight ∼21 hours), and 100% Lowicryl (overnight ∼24 hours). To ensure that there is no residual methanol, the samples were additionally incubated with fresh 100% Lowicryl for 1 hour. Then, samples were transferred into flat mold with 100% Lowicryl, incubated for additional 24 hours, and polymerized with UV light at -45°C for 27 hours, at 0°C for additional 40 hours, and at 20°C for 48 hours. The resin blocks were trimmed (Leica EM TRIM2) to reach a desired sample distance of 20-50µm from the upper surface of the block. Samples were then sputter coated with 25 nm of platinum (Electron Microscopy Science, EMS150T ES) and serial sectioned with a focused ion beam at a 20 nm step size and imaged in “Slice and View” mode with a backscattered electron detector using the FEI Helios 660 Nanolab system.

### Auditory Brainstem Responses (ABRs) and Distortion Product Oto-Acoustic **Emission (DPOAE)**

Hearing thresholds of *Cib2^R186W^* mutant mice were evaluated by recording ABR and DPOAE. All ABR recordings, including broadband clicks and tone-burst stimuli at three frequencies (8, 16, and 32 kHz), were performed using an auditory-evoked potential RZ6-based auditory workstation (Tucker-Davis Technologies) with high frequency transducer RA4PA Medusa PreAmps. Maximum sound intensity tested was 100 dB SPL. TDT system III hardware and BioSigRZ software (Tucker Davis Technology) were used for stimulus presentation and response averaging. DPOAEs were recorded using an acoustic probe (ER-10C, Etymotic Research) and DP2000 system (version 3.0, Starkey Laboratory). To measure DPOAE levels (2f1-f2), two primary tones, with a frequency ratio of f2/f1 = 1.2, were presented at intensity levels f1 = 65 dB SPL and f2 = 55 dB SPL, with f2 varied between 8–16 kHz (in one-eighth octave steps).

### Structural analysis of CIB2-TMC1/2

An Alphafold model of mouse CIB2 was obtained from the AlphaFold Protein Structure Database ^60^. An Alphafold multimer model ^61^ of the CIB2/TMC1/CIB2/TMC2 protein complex was generated using the AlphaFold Colab server without template constraint (https://colab.research.google.com/github/deepmind/alphafold/blob/main/notebooks/AlphaFold.ipynb). Protein/protein interfaces were analyzed using PDBePISA (https://www.ebi.ac.uk/pdbe/pisa/) ^62^. Structural models were analyzed, and figures prepared using PyMOL (The PyMOL Molecular Graphics System, Version 2.4.1 Schrödinger, LLC.)

## SUPPLEMENTAL FIGURES

**Supplemental Figure 1:**
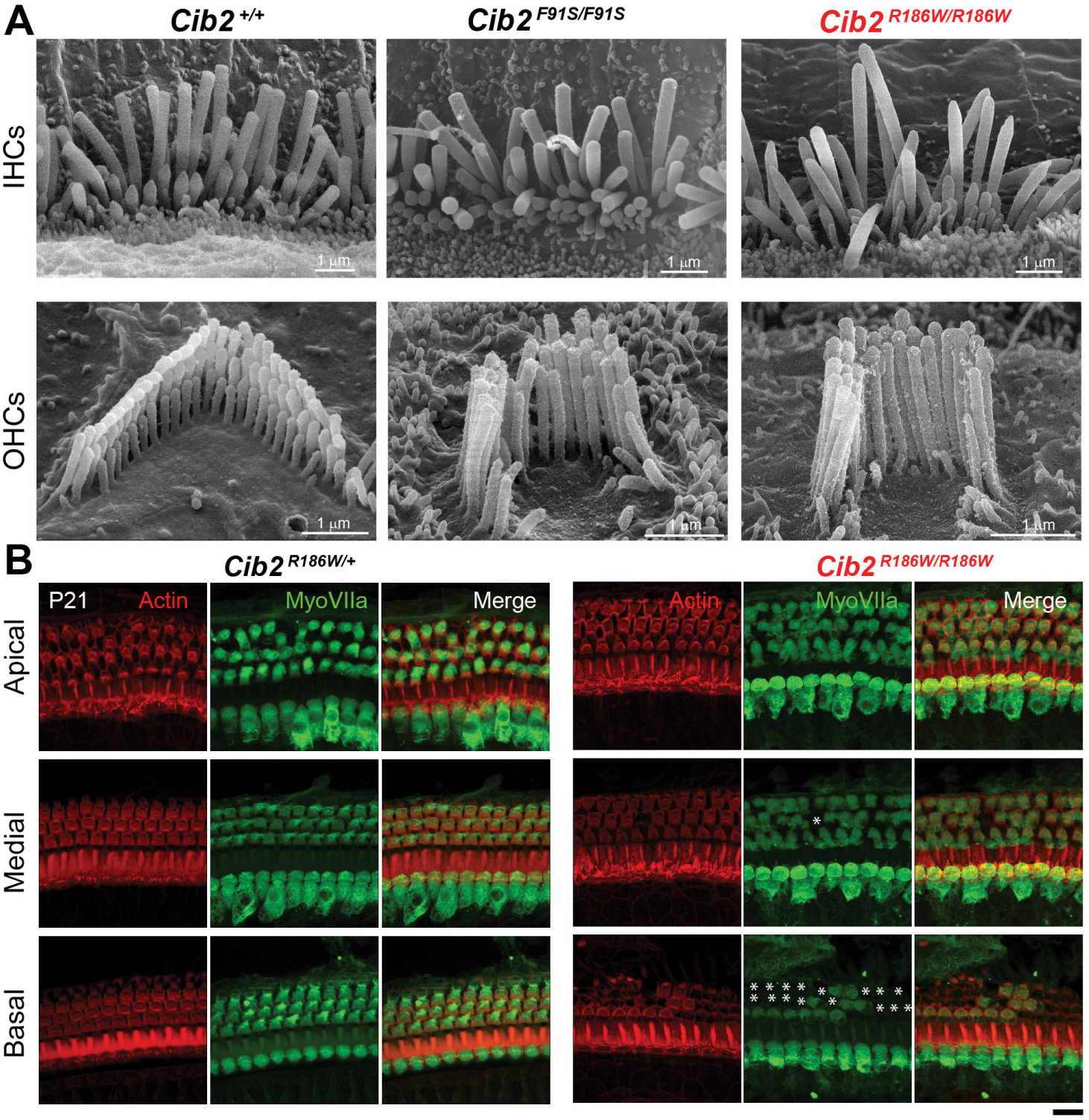
Similar to other deafness associated CIB2 variants, R186W variant causes stereocilia bundle abnormalities and progressive loss of OHCs. (A) Scanning electron microscopy (SEM) images of IHCs (top) and OHCs (bottom) in acutely isolated organ of Corti explants from wild-type (left), *Cib2^F91S/F91S^* (middle), and *Cib2^R186W^*^/*R186W*^ (right) mice at P17 showing similar hair bundle phenotypes across *Cib2* mutants. All cells were located approximately in the middle of the cochlea. (B) Maximum intensity projections of confocal Z-stacks of the apical (top), medial (middle), and basal (bottom) turns of *Cib2^R186W/+^*and *Cib2^R186W/R186W^* organs of Corti immunostained with myosin VIIa antibody (green) at P21 and counterstained with phalloidin (red). Asterisks indicate missing hair cells. Scale bar: 10 μm.

**Supplemental Figure 2:**
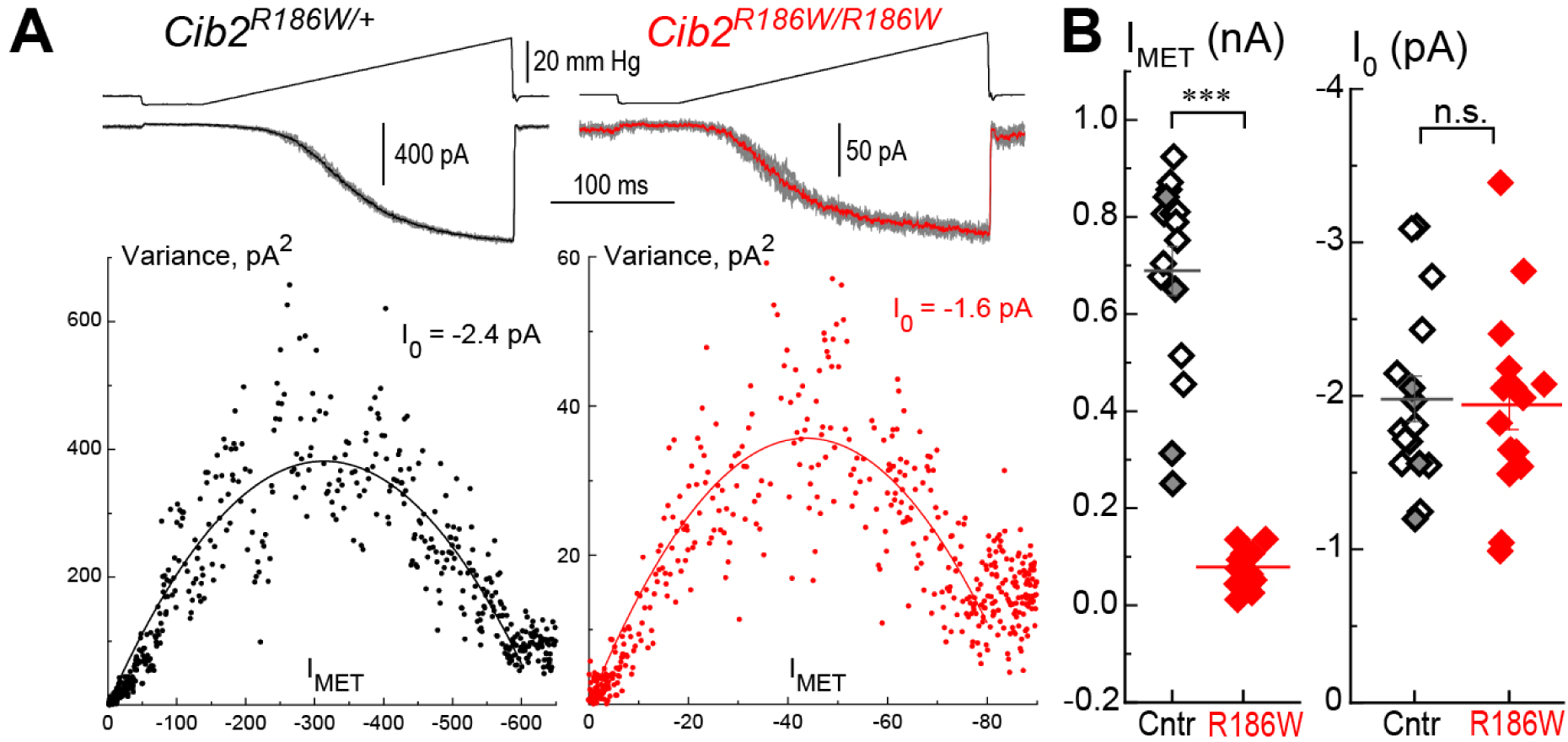
R186W variant does not affect single MET channel conductance in OHCs at P4-P6. (A) Individual MET current responses (middle, light gray or red) evoked by five identical fluid-jet ramp stimuli (top) with the mean current superimposed (black or red) in *Cib2^R186W/+^*(left) and *Cib2^R186W/R186W^* (right) OHCs. Bottom panels show the relationships between MET current variance and amplitude that was fit with a parabolic equation (see Methods) giving the apparent single-channel current of I_0_ = -2.4 pA and I_0_ = -1.6 pA for these *Cib2^R186W/+^* and *Cib2^R186W/R186W^* OHCs, correspondingly. (B) Maximal MET current at -90 mV (I_MET_, left) and the apparent single MET channel currents (I_0_, right) in control and *Cib2^R186W/R186W^* OHCs. Data from the same individual cells are shown on both graphs as well as mean±SE values. Asterisks indicate statistical significance of the differences: ***, P<0.001. Age of the cells: P4-P6, all of them were located approximately in the middle of the apical turn of the cochlea. Number of cells/mice: control, n=15/9 (*Cib2^+/+^*, n=7/3; *Cib2^R186W/+^*, n=8/6); *Cib2^R186W/R186W^*, n=15/7.

**Supplemental Figure 3:**
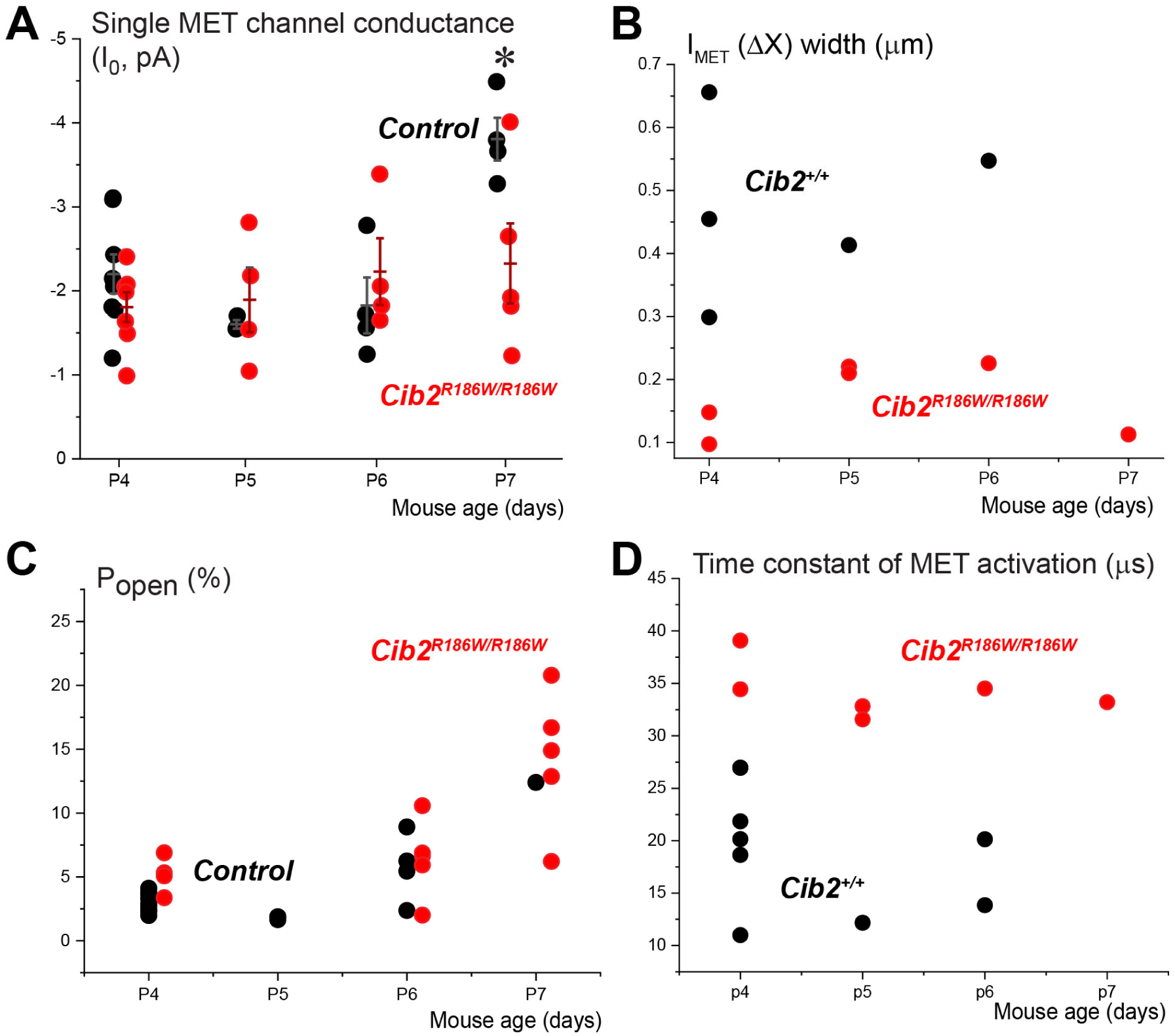
The abnormalities of MET currents in *Cib2^R186W/R186W^* OHCs are largely independent of the mouse age between P4 and P6. (A) Single MET channel conductance estimated by non-stationary fluctuation analysis (see Supplemental Fig. 2) at different ages. Data from individual wildtype (black) and *Cib2^R186W/R186W^* (red) OHCs are shown with mean±SE. Asterisks indicate significance: *, p<0.05 (Student *t*-test of independent variables). X-coordinate (age) was slightly shifted between the groups to avoid overlap. (B) Widths of the current-displacement curves (I-X) determined as in Fig. 3C-D at different ages. (C) Resting open MET channel probability (P_open_) at holding potential of -90 mV at different ages in individual wildtype (black) and *Cib2^R186W/R186W^*(red) OHCs. (B) No apparent changes of the time constant of MET channels activation with age. All the electrophysiology experiments were performed on mice between P4-P7. Each data point represents one cell and was calculated as an average of the time constants at bundle deflections of 0.1-0.8 µm (see Fig. 5D).

**Supplemental Figure 4:**
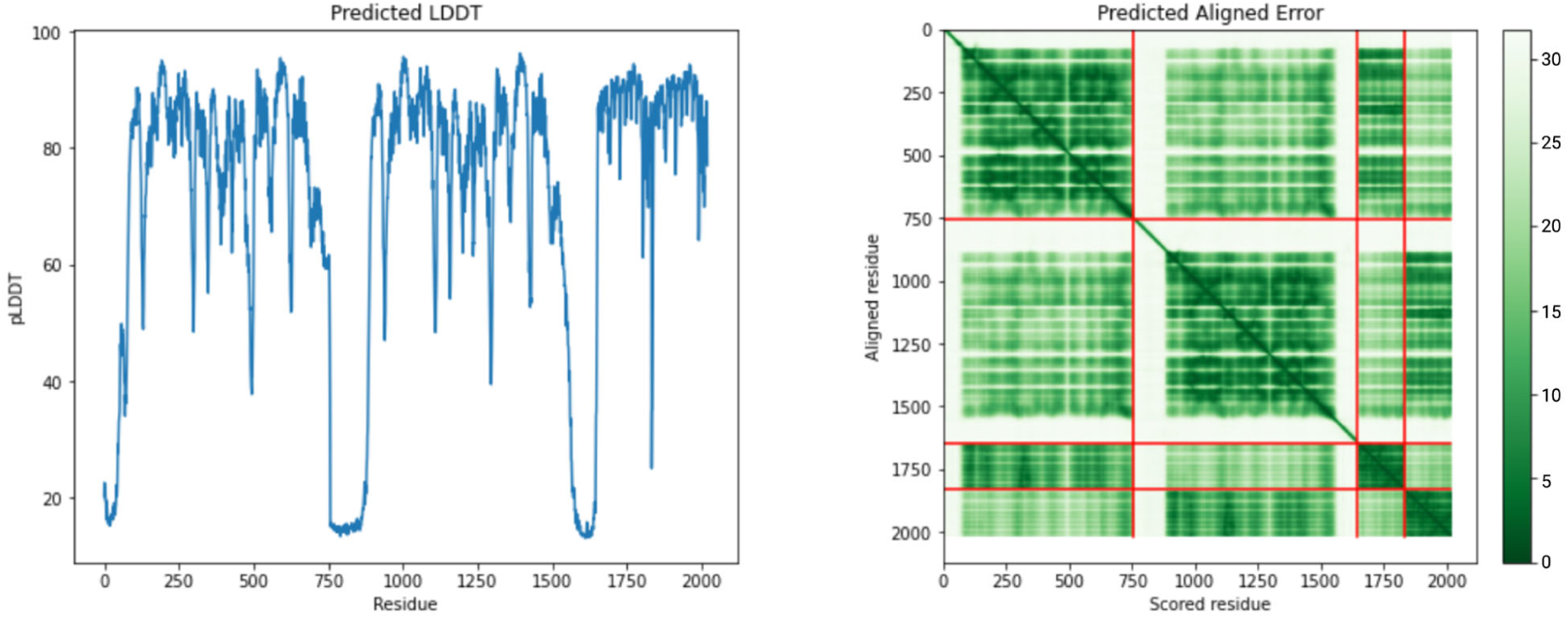
Predicted Local–Distance Difference Test (pLDDT) and Predicted Aligned Error (PAE) plots for the AlphaFold2 multimer prediction for the CIB2-TMC1/2 complex (Fig. 3G).

**Supplemental Figure 5:**
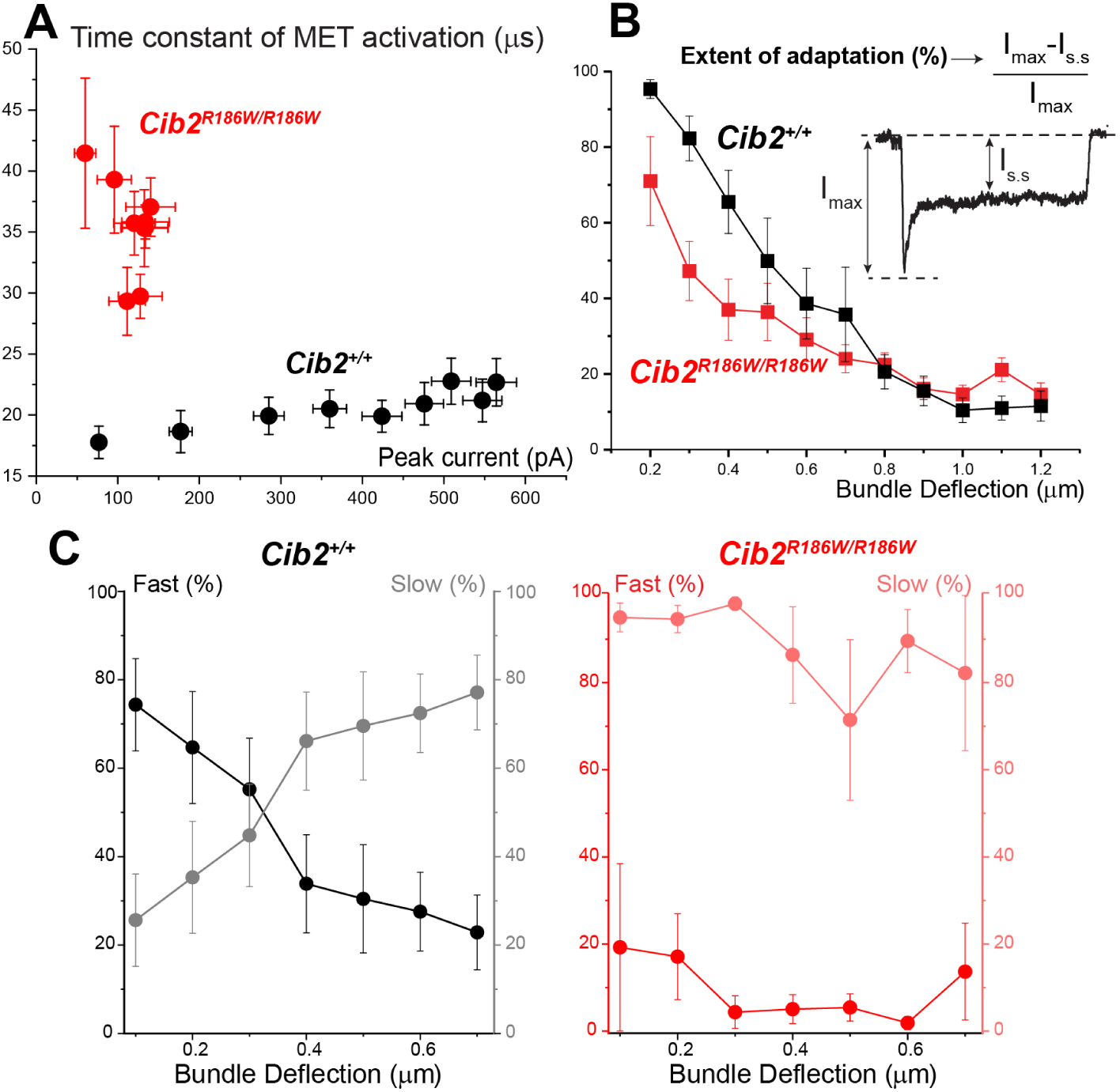
R186W variant of CIB2 increases the time constant of MET channel activation and disrupts fast adaptation. (A) Relationship between the time constants of MET channel activation and the amplitude of the MET current in wild type (black) and *Cib2^R186W/R186W^*(red) OHCs. Each data point represents the average of the time constants (Y-axis) and the peak MET currents (X-axis) at different step bundle deflections between 0.1 and 0.8 µm (see Fig. 5D). The number of cells/mice: *Cib2^+/+^*, n=16/12; *Cib2^R186W/R186W^*, n=6/6. (B) Relationship between the extent of adaptation (calculated as a ratio between the peak and the steady-state MET currents, see inset) and the hair bundle displacement. Means±SE are shown. The number of cells/mice: *Cib2^+/+^*, n=7/4; *Cib2^R186W/R186W^*, n=11/6. (C) Relative amplitudes of fast and slow adaptation determined from a double exponential fit of the MET currents in the wild type (left) and *Cib2^R186W/R186W^*OHCs (right). In many *Cib2^R186W/R186W^* OHCs, double exponential fit was not possible and, therefore, the amplitude of fast adaptation was set to zero.

